## Supplementary materials for "A model for learning based on the joint estimation of stochasticity and volatility"

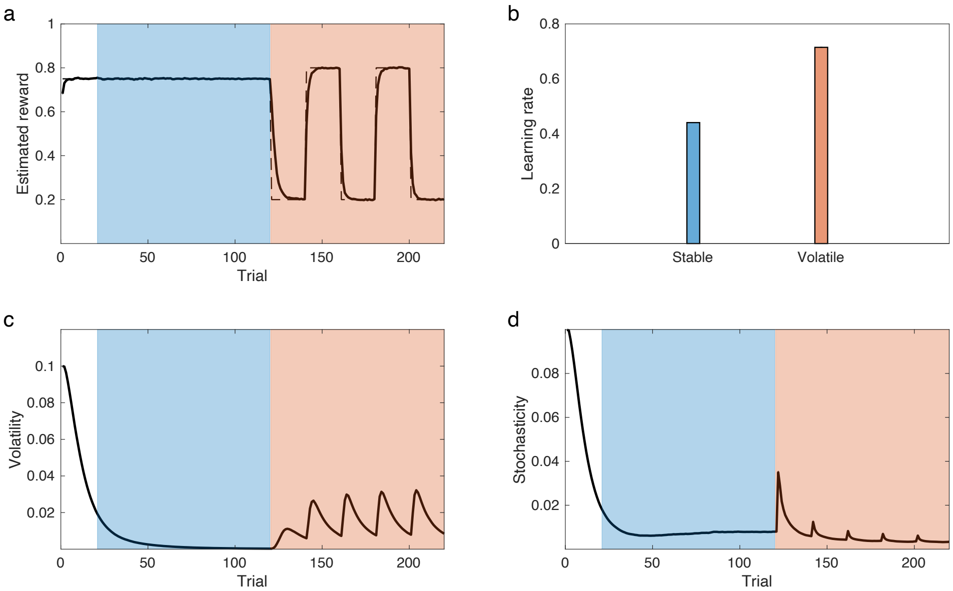


Supplementary Fig 1. The model elevates its learning rate in volatile environments. a) Simulations of our model in the volatility learning paradigm, in which subjects undergo stable (bluish) and volatile blocks (orangish) of learning. Dashed and solid line show true reward and estimated reward by the model, respectively. b) learning rate is larger in the volatile block compared with the stable one, similar to those reported in humans. This is because the volatility term (c) increases more than the stochasticity term (d) in the volatile condition. Errorbars reflect standard error of the mean over 1000 simulations and are too small to be visible.

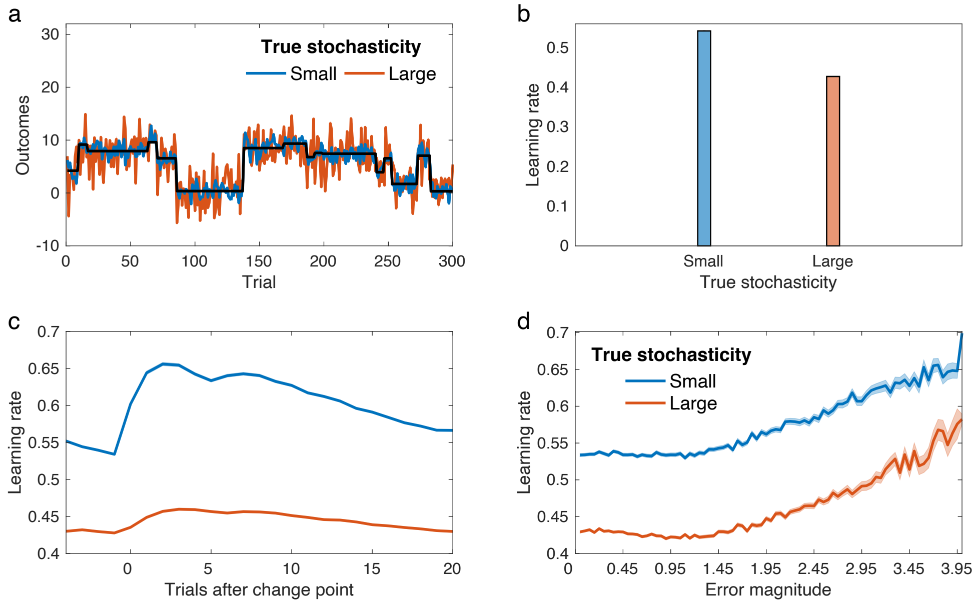


Supplementary Fig 2. The model reduces the learning rate in stochastic environments. a) The prediction task by Nassar and colleagues[1], in which the participant makes a new prediction of outcome on every trial. Outcomes are generated based on a true reward rate, which undergoes occasional jumps, plus small or large amount of noise (i.e. true stochasticity). b) Behavior of the model in this task: Increases in the noise level is analogous to increases in stochasticity, which decreases the learning rate. The model also explains aspects of empirical data (Nassar et al.[1]) that are quite independent of stochasticity and are more closely related to jumps (c-d). c) Learning rate increases following switches in the task for both types of noise, although this effect is stronger for smaller noise level. d) The model learning rate also increases by increase in absolute error magnitude (divided by the value of true noise). Errorbars reflect standard error of the mean over 1000 simulations and are too small to be visible.

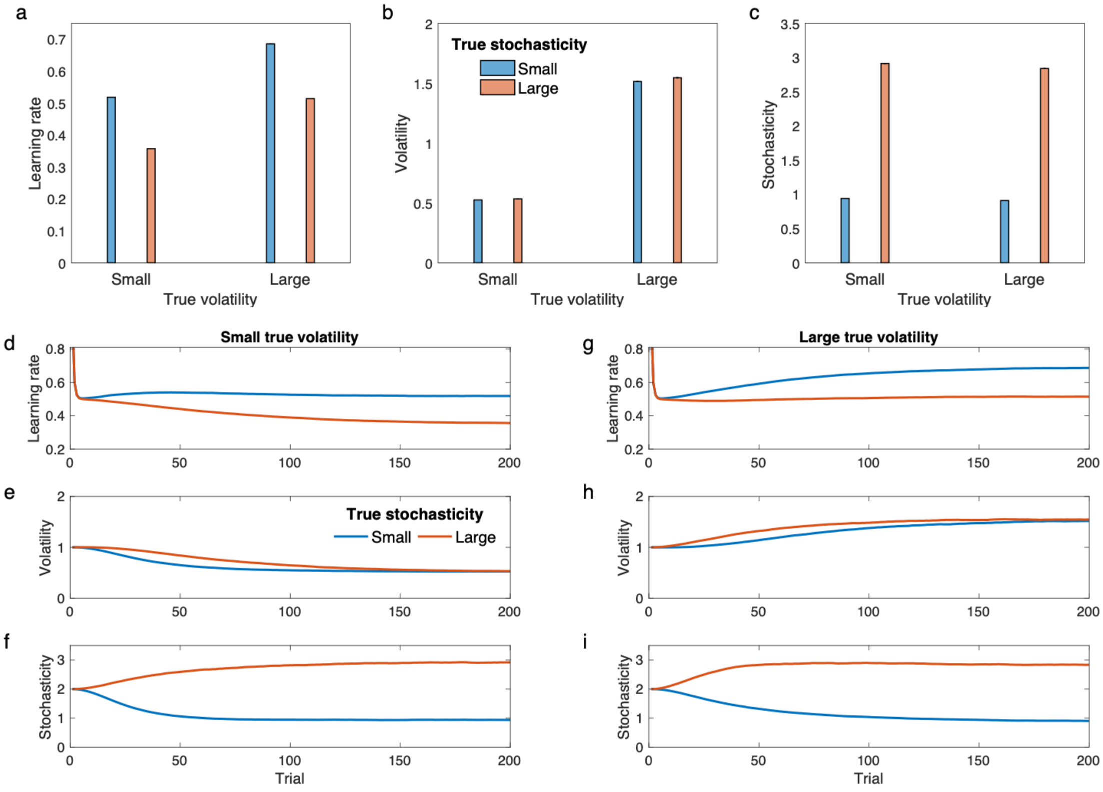


Supplementary Fig 3. Simulations in the same setting of Fig. 2 for an alternative model in which the generative process for volatility and stochasticity is assumed to be Gaussian (see Methods). The same particle filter has been used for inference. These results are very similar to the original results reported in Fig. 2 indicating that the model is not sensitive to the generative process as long as the same inference algorithm has been employed. Model parameters were assumed to be $\sigma_v^2 = \sigma_s^2 = 0.1$. Errorbars reflect standard error of the mean over 10000 simulations and are too small to be visible.

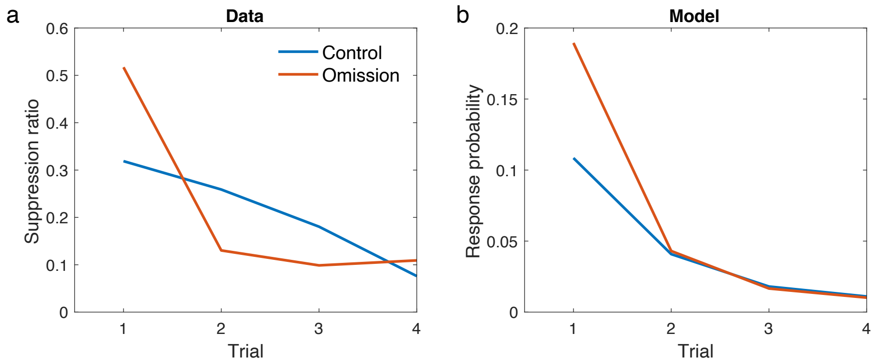


Supplementary Fig 4. Suppression ratio reported by Hall and Pearce (a) and the median response probability by the model in the retraining phase (b). The omission group shows faster decrease, consistent with empirical data. Suppression ratio has been defined as the ratio of response in the 90 seconds window following presentation of the cue divided by its sum with the response rate in the preceding window of 90 seconds. Data in a adapted from Hall and Pearce[2].

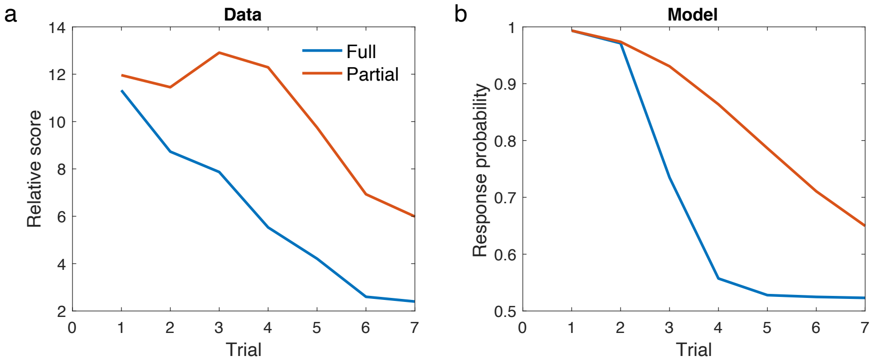


Supplementary Fig 5. Empirical data for the partial reinforcement effect experiment (a) and the median response probability by the model in the retraining phase (b). Empirical data of the retraining phase were reported by Haselgrove et al.[3], in which the relative score has been calculated by subtracting the duration of magazine activity during the pre-CS period from duration of the magazine activity during the CS period. The average relative score across two sessions of extinction has been plotted.

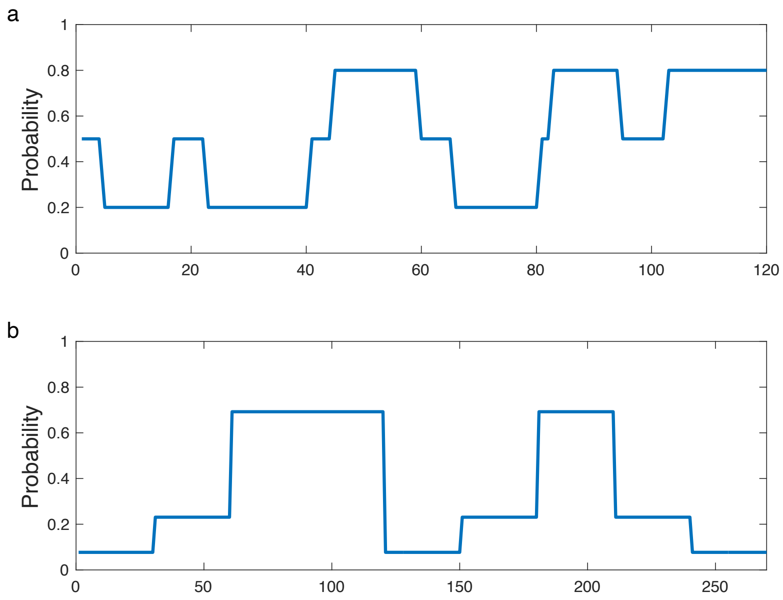


Supplementary Fig 6. Probability sequences used for simulations presented in Fig. 5f (a) and 5h (b). For simulating the second task (b), the actual change-points were chosen based on a normal distribution with the mean 30 (from the previous change point) and variance 1, similar to the original experiment.

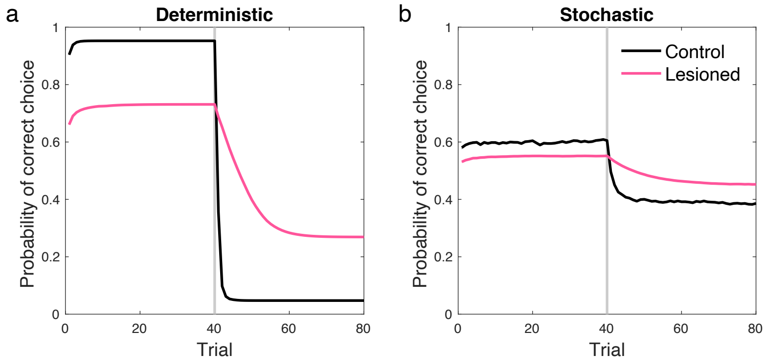


Supplementary Fig. 7. Probability of correct choice for simulation results reported in Fig. 8, which is well matched by empirical data reported by Costa et al[4].

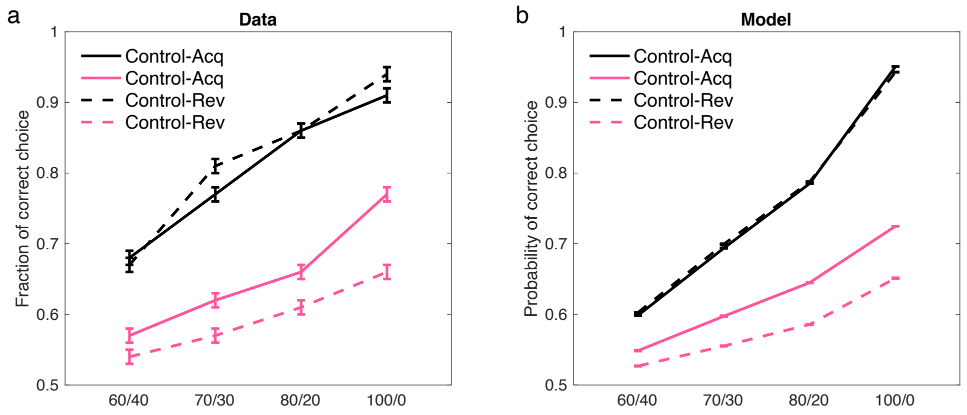


Supplementary Fig. 8. Simulation of the model for all probabilistic schedules tested by Costa et al.[4]. a-b) empirical data and model simulations are plotted. Errorbars reflect standard error of the mean (over 1000 simulations in (b)).

| Group | Data | Model |
| --- | --- | --- |
| **Control-consistent** | 53.9 | 56.18 |
| **Control-shift** | 55.6 | 56.22 |
| **Lesioned-consistent** | 59.9 | 56.21 |
| **Lesioned-shift** | 52.2 | 56.21 |

Supplementary Table 1. Reported data (time spent in the food cup%) of phase 1 of the experiment by Holland and Gallagher[5] (Fig. 7) following presentation of the second cue. No significant difference was found between control and lesioned animals suggesting that lesioned animals were able to learn efficiently. The model shows the same behavior (percentage of food response is reported, which is calculated based on a softmax with the decision noise of 0.5).

### References


1. Nassar, M. R., Wilson, R. C., Heasly, B. & Gold, J. I. An approximately Bayesian delta-rule model explains the dynamics of belief updating in a changing environment. *J. Neurosci.* **30**, 12366–12378 (2010).
2. Hall, G. & Pearce, J. M. Restoring the Associability of a Pre-Exposed CS by a Surprising Event. *Q. J. Exp. Psychol. Sect. B* **34**, 127–140 (1982).
3. Haselgrove, M., Aydin, A. & Pearce, J. M. A partial reinforcement extinction effect despite equal rates of reinforcement during Pavlovian conditioning. *J. Exp. Psychol. Anim. Behav. Process.* **30**, 240–250 (2004).
4. Costa, V. D., Dal Monte, O., Lucas, D. R., Murray, E. A. & Averbeck, B. B. Amygdala and Ventral Striatum Make Distinct Contributions to Reinforcement Learning. *Neuron* **92**, 505–517 (2016).
5. Holland, P. C. & Gallagher, M. Amygdala central nucleus lesions disrupt increments, but not decrements, in conditioned stimulus processing. *Behav. Neurosci.* **107**, 246–253 (1993).
